# Mutation of *GmDMP* genes triggers haploid induction in soybean

**DOI:** 10.1101/2024.03.24.585499

**Authors:** Yu Zhong, Mingliang Yang, Dehe Cheng, Jinchu Liu, Qi Han, Chunyan Liu, Xiaolong Qi, Tongzheng Yan, Lei Teng, Chang Xv, Jingjing Hou, Lianjun Sun, Chenxu Liu, Qingshan Chen, Shaojiang Chen

**Affiliations:** National Maize Improvement Center of China, Key Laboratory of Crop Heterosis and Utilization/Engineering Research Center for Maize Breeding, Ministry of Education, College of Agronomy and Biotechnology, China Agricultural University, Beijing 100193, China; College of Agronomy and Biotechnology, China Agricultural University, Beijing 100193, China; College of Agriculture, Northeast Agricultural University, Harbin 150030, China; College of Horticulture, China Agricultural University, Beijing, 100193, China

**Keywords:** DH, soybean, haploid induction gene, spontaneous diploidization

## Abstract

The development of homozygous lines is a key step in plant breeding and production. Generally, homozygous lines can be obtained through traditional time-consuming way of several generations selfing or through a way of doubled haploid (DH) technology, which has obvious advantages to accelerate breeding. However, no effective haploid production system so far has been established in soybean. Here we show that mutations of the soybean *GmDMP1* and *GmDMP2* genes can be used to induce haploid with an average haploid induction rate of 0.61%. We also found that 22.9% of soybean haploids can produce seeds through spontaneous chromosome doubling. Those findings laid a solid foundation for establishing DH technology in soybean, which will accelerate soybean breeding.

## Introduction

Soybean (*Glycine max* [L.] Merr.) is the most economically important legume in the world, providing more than 25% of the protein for human consumption and livestock feed^[1]^. The major goals of soybean improvement are to increase yield and expand adaptability. Even though soybean breeding has benefited greatly from advances in genomics and genetics, the development of homozygous lines in soybean still relies on conventional breeding methods^[2]^. These approaches generally require selfing for more than eight generations, which is costly and time-consuming^[3]^. Alternatively, doubled haploid (DH) technology can be used to obtain homozygous lines in only two generations^[4]^.

Haploids can be induced *in vitro* from the culture of gametophyte cells or *in vivo* by interspecific crosses or by intraspecific crosses with haploid inducer lines or manipulation of *CENH3* centromeric histone proteins^[5]^. Although there are several haploid induction (HI) systems available, HI in soybean has been a long-standing challenge worldwide^[6]^. Despite a lot of efforts being made to develop an *in vitro* HI system in soybean over the past six decades^[6-9]^, there is still no efficient method available for the production of soybean haploid due to genotype-dependent and methodological complexity.

Currently, the most efficient method for haploid production is *in vivo* HI triggered by haploid inducer line, which has been used successfully and extensively in maize breeding. The identification of the causal genes opens up the possibility for the development of a similar HI system in other crops^[10, 11]^. Based on the *ZmPLA1*/*MATL*/*NLD* gene, *in vivo* haploid inducer lines have been created in rice, wheat, and other cereals^[4]^. The utility of *dmp* mutants for HI in dicots was recently demonstrated in ten species from five different families^[3, 12-16]^. Here, we show that mutation of *GmDMP* genes also triggers haploid induction in soybean.

## Results

We previously identified two *DMP* genes (*GmDMP1*/*GLYMA_18G097400*; *GmDMP2*/*GLYMA_18G098300*) in soybean using a three-step *DMP* selection strategy^[15]^. These genes are specifically expressed in flower buds and share 61% amino acid sequence identity with *ZmDMP*^[15]^. Furthermore, *GmDMP1* and *GmDMP2* exhibit a high degree of amino acid sequence identity (97%) and are closely linked on the same chromosome (∼140 kb), indicating that they are functionally redundant and more likely to be inherited together during recombination. Therefore, a CRISPR-Cas9 mutagenesis construct comprising four sgRNAs was designed to generate designed to simultaneously generate double mutants in both *GmDMP1* and *GmDMP2* in the Williams 82 background. To facilitate haploid identification, both the *35S*:*GFP* and the FAST-Red maker were included in the construct (Figure 1a). Among 21 T_0_ transgenic events, we identified two lines that carried the mutation in both *DMP* genes. In the T2 generation, we obtained two homozygous *gmdmp* mutants, M1 and M2 (Figure 1b). No obvious abnormal vegetative growth phenotypes were observed in these *gmdmp* mutants, but pleiotropic seed phenotypes were observed in self-pollinated progeny (Figure 1c). These data are in line with previous reports in *arabidopsis* and tomato *dmp* mutants^[12, 14]^.

**Figure 1.**
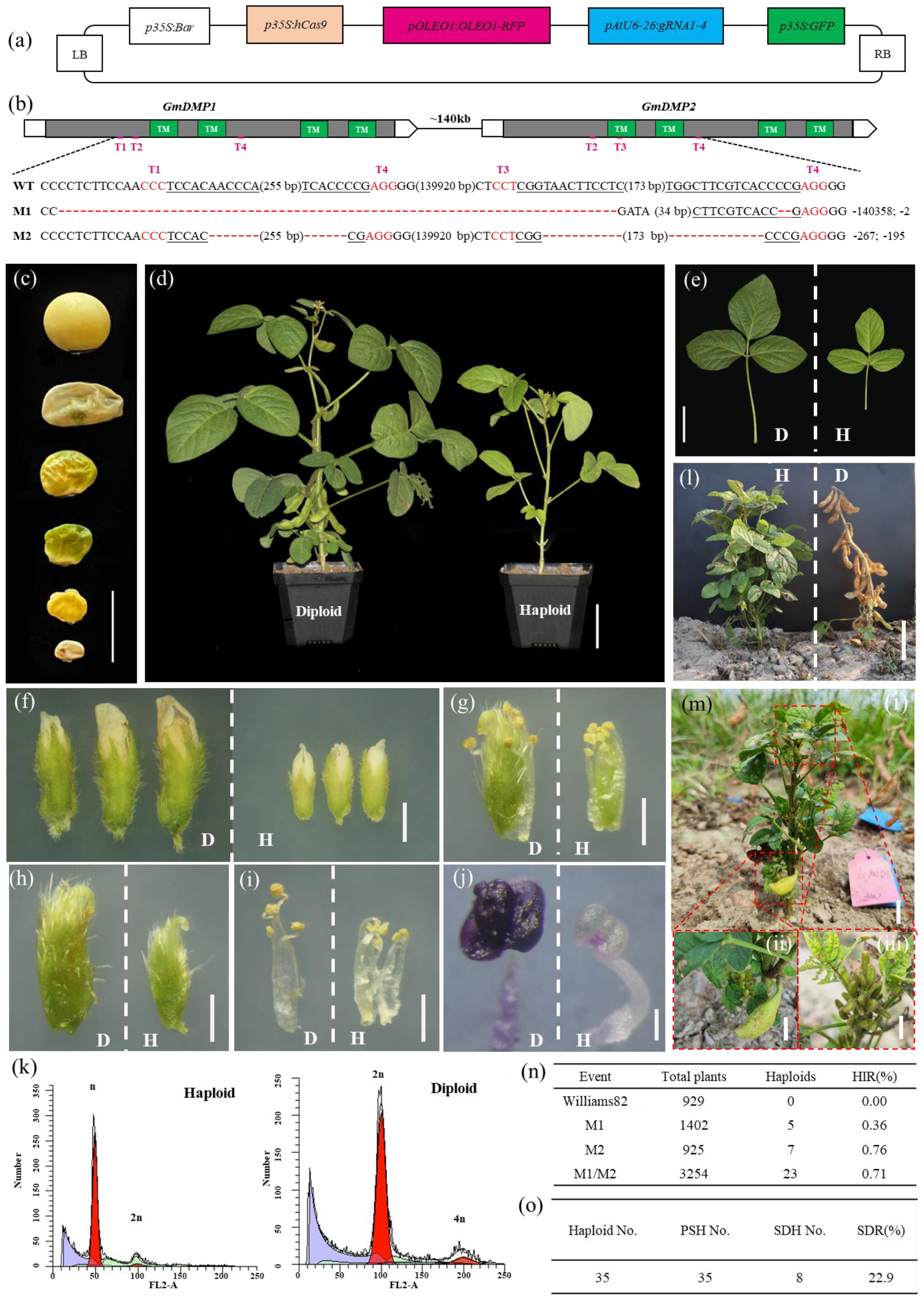
Mutation of *GmDMP* genes triggers haploid induction in soybean. (a)The CRISPR/Cas9 mutagenesis vector comprising four sgRNAs targeting *GmDMP*. (b)Schematic representation of *GmDMP1* and *GmDMP2* gene structures and genome editing experimental design. Filled blocks indicate the coding region. Green blocks correspond to the regions encoding the four predicted transmembrane domains (TMs). Red lines indicate the four regions (T1–4) targeted by sgRNAs. The relevant sequences from the wild-type (WT) and mutant alleles are shown below the gene structure schematics. (c) Pleiotropic seed phenotypes in self-pollinated *dmp* mutants. (d-i) Phenotypes of plants (d), leaves (e), flower buds (f), and dissected flower buds (g, h, i) of haploid and diploid plants. (j) Haploid (H) and diploid (D) anthers after staining with Alexander staining solution. (k) Haploid and diploid plants at maturity stage. (l) Haploid plant (i) set smaller pods (ii, top of the plant) and bulging pod (iii, bottom of the plant). (m) Flow cytometry verification of the ploidy of a putative haploid and a diploid control. The x-axis represents the signal peak for the nucleus and the y-axis represents the number of nuclei. (n) Haploid induction rate (HIR) of *gmdmp* mutants. (o) Spontaneous diploidization in soybean haploids. No., number; PSH, pod set haploid; SDH, spontaneous diploidization haploid; SDR, spontaneous diploidization rate. Scale bars: 1 cm (c, l), 5 cm (d), 4 cm (e), 2 mm (f), 1 mm (g, h, i), 0.1 mm (j), 6cm (k).

To investigate whether *dmp* mutants can induce haploid in soybean, we sowed selfed seeds from T_3_ progenies of M1, M2 and M1 × M2 mutants. Without segregating molecular markers, we identified putative haploid plants based on their phenotype and then assessed for ploidy level by flow cytometry^[11, 12]^. In the *gmdmp* mutant progeny, we found that some plants were stunted in growth and development, with smaller leaves and sterile floral organs (Figure 1d-j). In addition, these plants retained green leaves even when others were ready for harvest (Figure 1k, l). Most of these plants were verified to be true haploids by flow cytometry (Figure 1m). In the end, five haploids (0.36%) were found in 1,402 M1 mutant plants, seven haploids (0.76%) in 925 M2 mutant plants, 23 haploids (0.71%) in 3,254 M1 × M2 mutant plants, whereas no haploids were found in 929 wild-type plants (Figure 1n). The results indicate that mutation of the *GmDMP* genes induces *in vivo* haploid production in soybean.

Haploid plants are sterile and require chromosome doubling to generate DH lines. Chromosome doubling is usually induced chemically, but spontaneous doubling of haploid embryos *in vitro* is also commonly observed. Spontaneous diploidization was observed from *dmp*-triggered soybean haploid plants, as evidenced by the production of pods and viable seeds (Figure 1m). All 35 haploids set smaller pods at the top of the plants, of which eight (22.9%) eventually produced viable seeds (Figure 1o). This result indicates that soybean haploids might have a high potential for spontaneous chromosome doubling, which can give rise to a higher proportion of spontaneous DH lines.

## Discussion

In summary, our study demonstrates for the first time that mutation of *GmDMP* genes can trigger *in vivo* haploids in soybean. In addition, the study found the phenomenon of spontaneous diploidization in soybean haploids, indicating that it is possible to produce DH lines naturally. These findings laid a solid foundation for establishing DH technology in soybean, which will greatly accelerate soybean breeding.

## Materials and methods

### Vector construction

Primers used to amplify and sequence the *DMP* alleles were designed based on the *GmDMP1* and *GmDMP2* sequences downloaded from the National Center for Biotechnology Information. The U6-26 promoter was used to drive expression of all four sgRNAs. The *bar* gene driven by the cauliflower mosaic virus (CaMV) *35S* promoter was used for positive callus selection during transformation. FAST-Red cassette and *GFP* driven by *35S* promoter were used for the identification of haploid seeds and seedlings, respectively. All these cassettes, together with human codon-optimized Cas-9 driven by a 35S promoter, were introduced simultaneously in one step into pISCL4723 through the Golden Gate cloning method^[17]^. Supplementary Table 1 lists the primers for vector construction.

### Plant materials

The soybean cultivar Williams 82 was used as a receptor line for *Agrobacterium*-mediated transformation as previously described.

### Screening for *gmdmp* mutants

The *gmdmp* mutants were identified in T0 transgenic lines by PCR amplification and sequencing of the *DMP* locus. PCR products containing multiple amplification products from the same locus were further amplified with KOD FX (Toyobo) and cloned into the pEASY vector (pEASY-Blunt Zero Cloning Kit; TransGen Biotech) and then to Escherichia coli. At least six independent E. coli colonies per corresponding PCR reaction were selected and sequenced. All sequences were aligned to the *GmDMP* wild-type alleles with SnapGene software. The primers used for *gmdmp* mutants genotyping are listed in Supplementary Table 1.

### Haploid identification

Putative haploids from selfed *gmdmp* mutant lines were initially identified by their phenotype (small organs, stunted in growth and development) and then confirmed by flow cytometry.

### Flow cytometry

Fresh leaves (0.5 g) from each sample were chopped with a razor blade in 2 mL lysis buffer as previously described and filtered through an 80 μm nylon filter. Nuclei were collected by centrifugation at 1000 r.p.m. for 5 minutes at 4 °C and stained with propidium iodide in the dark for 20 min. The ploidy level of each sample was analyzed with a BD FACSCalibur Flow Cytometer and BD CellQuest Pro software. Diploid wild-type soybean was used as a control and the position of its first signal peak was set at∼100 (FL2-A value). The samples with the first signal peak at∼25 (FL2-A value) were deemed to be haploids.

## Acknowledgements

This work was supported by the National Natural Science Foundation of China (32201836, 32341036), the National Key Research and Development Program of China (2022YFD1200802, 2022YFF1003204), Beijing Nova Program (2023067) and the Modern Maize Industry Technology System (CARS-02-05).

## Author contributions

Y.Z., C.L., Q.C. and S.C. conceived and designed the experiments. Y.Z., D.C. and M.Y. performed most of the experiments. J.L., Q.H., C.L., X.Q., T.Y., L.T., C.X., and J.H. performed some of the experiments. Y.Z., D.C., L.S., C.L., Q.C. and S.C. analyzed the data. Y.Z., D.C. and S.C. discussed and prepared the manuscript. All authors discussed the results and provided feedback on the manuscript.

---

**Supplementary Table 1.**
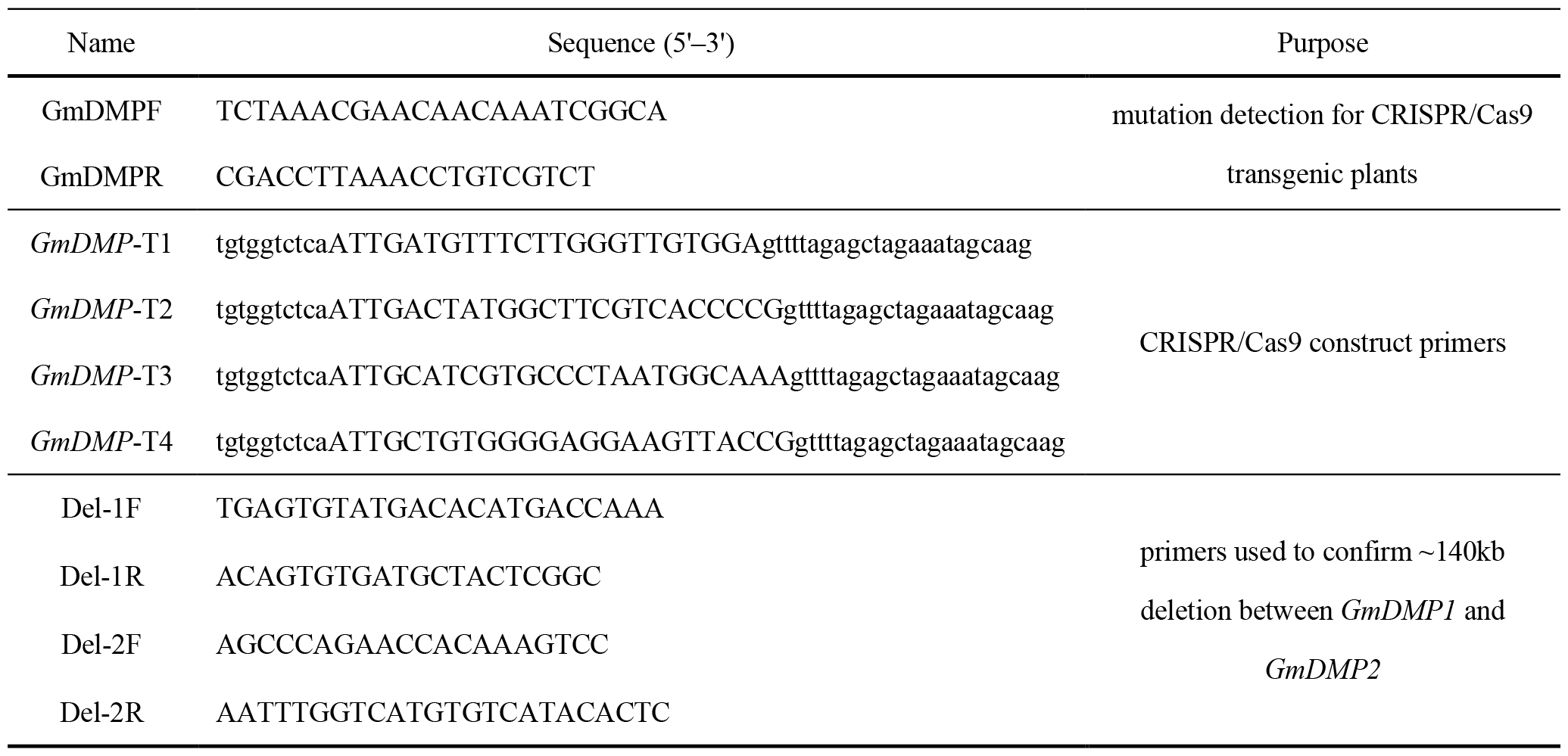
Primers used in this study.

